# Context matters: environmental microbiota of ice cream processing facilities affects the inhibitory performance of two lactic acid bacteria against *Listeria monocytogenes*

**DOI:** 10.1101/2022.10.04.510917

**Authors:** M. Laura Rolon, Tyler Chandross-Cohen, Kerry E. Kaylegian, Robert F Roberts, Jasna Kovac

**Author notes:** Corresponding author: Jasna Kovac.

## Abstract

Pathogenic *L. monocytogenes* may inhabit dairy processing environments, increasing the risk for cross-contamination of foods. Using biocontrol microorganisms that inhibit or outcompete *L. monocytogenes* to complement sanitation of dairy processing facilities may enhance the control of *L. monocytogenes*. However, it remains unknown whether the resident microbiota of dairy processing facilities affects the antilisterial activity of biocontrol strains. Here, two lactic acid bacteria (LAB) strains (*Enterococcus* PS01155 and PS01156) were tested for their biocontrol potential in the context of microbiomes collected from three ice cream processing facilities (A, B, and C). Antilisterial ability was assessed by co-culturing LABs with 8-*L. monocytogenes* strains in the presence of microbiota for 3 days at 15°C, followed by quantification of the most probable number of attached *L. monocytogenes. L. monocytogenes* concentration increased by 0.38±0.77 log_10_ MPN/sample in treatments containing microbiota from facility A, while it decreased by 0.99±1.13 and 2.54±0.84 log_10_ MPN/sample in treatments with microbiota from facilities B and C, respectively. The attachment of LAB to an abiotic surface was assessed by co-culturing LABs in with the microbiomes at 15°C for 3 days, followed by characterization of attached microbiota composition using amplicon sequencing. All samples containing microbiomes from facilities A and B had high relative abundance of *Pseudomonas*, while samples with facility C microbiome had high relative abundance of *Enterococcus*. Overall, we show that microbiota composition of ice cream processing facilities affected the antilisterial ability of LABs.

**IMPORTANCE:** Antilisterial lactic acid bacteria strains had been proposed as biological pathogen control agents for application in food processing environments. However, the effect of resident food processing environment microbiota on the performance on antilisterial lactic acid bacteria strains is poorly understood. Our study shows that the composition of the microbiota collected from ice cream processing facilities’ environmental surfaces can affect the attachment and inhibitory effect of lactic acid bacteria strains against *L. monocytogenes*. Further studies are therefore needed to evaluate whether individual microbial taxa affect antilisterial properties of lactic acid bacteria strains and to characterize the underlying mechanisms.

---

The foodborne pathogen *Listeria monocytogenes* causes listeriosis, a rare but deadly disease. In the United States, foodborne listeriosis is associated with a 94% hospitalization rate and a 15.9% death rate (Scallan et al., 2011). Listeriosis outbreaks have been linked to the consumption of dairy foods, including raw milk, cheese, and ice cream (Lundén et al., 2004). In 2015-2016, a multistate outbreak of listeriosis traced to ice cream products served in hospital settings (Rietberg et al., 2016) revealed that *L. monocytogenes* can survive in frozen dairy products (Salazar et al., 2020). While *L. monocytogenes* is effectively inactivated by pasteurization (Ceylan et al., 2017), it may be re-introduced to dairy products after heat treatment if present in the food processing environment (Melo et al., 2015). *L. monocytogenes* can form or attach to existing biofilms which allows it to persist in food manufacturing facilities (Melo et al., 2015). Biofilms can be formed by multiple species of microorganisms present in the food processing environments and provide a physical barrier to the antimicrobial action of sanitizers against *L. monocytogenes* (Mazaheri et al., 2021). It is therefore critical to effectively control the presence of *L. monocytogenes* in food processing environments to prevent contamination of products and maintain consumer trust.

Current strategies to manage *L. monocytogenes* in dairy processing environments include pathogen environmental monitoring (Beno et al., 2016) and standard cleaning and sanitization protocols (Marriott et al., 2018). Chemical treatments, such as the use of caustics and acids for cleaning, followed by the application of chlorine-based or quaternary ammonium-based sanitizers are the most frequent methods used in dairy processing facilities for cleaning and sanitizing (Marriott et al., 2018). However, the effectiveness of cleaning and sanitizing in dairy facilities may be compromised in difficult-to-clean areas (Carpentier & Cerf, 2011). Biocontrol bacteria (e.g., strains that inhibit or outcompete *L. monocytogenes*), can be intentionally added into sanitizing protocols to enhance the control of *L. monocytogenes* in food processing environments (Camargo et al., 2018).

Lactic acid bacteria, including *Enterococcus* spp., may inhibit *L. monocytogenes* through the production of organic acids, hydrogen peroxides, catalases, and bacteriocins (Camargo et al., 2018). Further, they can prevent the attachment of *L. monocytogenes* to environmental surfaces by competition for available nutrients and space (Camargo et al., 2018). For example, two lactic acid bacteria strains with antilisterial properties originally identified as *Enterococcus durans* 152 and *Lactococcus lactis* subsp. *Lactis* C-1-92 (Zhao et al., 2004), and later re-identified as *Enterococcus faecium* PS01155 and *Enterococcus lactis* PS01156 (Sinclair et al., 2022) were able to reduce the concentration of *Listeria* spp. by 2-4 log CFU/cm^2^ in floor drains of a poultry production facility (Zhao et al., 2006). However, it was not assessed whether the resident microbiota of drains could have affected the antilisterial action of the lactic acid bacteria applied as biocontrols. Microorganisms that are introduced as biological controls to new environments will interact with the resident microbiota and compete for space and resources. Because of the diversity of organisms that reside in dairy processing environments (Bokulich & Mills, 2013; Calasso et al., 2016; Dzieciol et al., 2016; Falardeau et al., 2019; Guzzon et al., 2017; Johnson et al., 2021; Schön et al., 2016; Stellato et al., 2015), it remains unknown whether and to what extent the use of specific biocontrols will be effective in controlling *L. monocytogenes* within a specific dairy processing facility.

In this study, we seek to assess the antilisterial ability of the lactic acid bacteria strains isolated by Zhao et al., (2004) in the context of environmental microbiota of ice cream processing facilities using an *in vitro* model system. We specifically aimed to (i) assess whether the two lactic acid bacteria strains can inhibit the growth of *L. monocytogenes* strains isolated from dairy processing environments, (ii) determine whether two lactic acid bacteria strains can effectively inhibit *L. monocytogenes* in the presence of environmental microbiomes collected from ice cream processing facilities, (iii) determine whether two lactic acid bacteria strains can attach to an abiotic surface in the presence of environmental microbiomes of ice cream processing facilities, and (iv) determine whether the presence of *Pseudomonas* spp. influence the antilisterial activity of two lactic acid bacteria.

## RESULTS

### The antilisterial ability of *Enterococcus faecium* PS01155 and *Enterococcus lactis* PS01156 was dependent on temperature, *L. monocytogenes* strain and concentration, and microbiome context

Two lactic acid bacteria strains of the genus *Enterococcus* (PS01155 and PS01156), isolated by (Zhao et al., 2004), were selected due to their reported ability to inhibit *L. monocytogenes* from drains in a poultry processing facility (Zhao et al., 2006). To determine whether these lactic acid bacteria strains could effectively inhibit *L. monocytogenes* strains isolated from dairy processing environments (Table 1) (Beno et al., 2016), we conducted a spot-inoculation assay on BHI agar plates against lawns of each *L. monocytogenes* strain at a concentration of ∼10^7^ or ∼10^8^ CFU/ml, incubated at 15, 20, 25, or 30°C. PS01155 and PS01156 were able to inhibit all tested strains of *L. monocytogenes* (Table 1) at 20, 25, and 30°C (Fig. 1A). On lawns of *L. monocytogenes* with a concentration of ∼10^7^ CFU/ml, PS01155 and PS01156 showed a significant increase in the inhibition of *L. monocytogenes* with an increase in temperature (p = 2.0 × 10^−16^). PS01155 did not inhibit any of the tested *L. monocytogenes* strains at 15°C. Further, it was significantly more inhibitory at 25°C compared to 20°C. Similarly, PS01156 was significantly more inhibitory to *L. monocytogenes* at 20, 25, and 30°C compared to 15°C. Similar trend was observed on lawns of *L. monocytogenes* with a concentration of ∼10^8^ CFU/ml, where PS01155 had a significantly higher inhibition at 25 and 30°C compared to at 20°C. PS01155 did not inhibit any of the tested *L. monocytogenes* strains at 15°C. In contrast to PS01155, the inhibitory activity of PS01156 was not temperature dependent. Furthermore, there was a significant difference in the antilisterial abilities of PS01155 and PS01156 influenced by the specific *L. monocytogenes* strain used to make the lawns (p < 2*10^−16^) (Fig. S1).

**TABLE 1.** Bacterial strains used for the study.

| **Species** | **Strain** | **Isolation source** | **PFGE pattern** | **Lineage** | **Reference** |
| --- | --- | --- | --- | --- | --- |
| *L. monocytogenes* | FSL A5-0335 / PS00975 | Dairy processing facility, swab, processing, drain | CU-172,425 | II | (Beno et al., 2016) |
| *L. monocytogenes* | FSL A5-0151 / PS00958 | Dairy processing facility, swab, aging room doorway | CU-311,37 | I | (Beno et al., 2016) |
| *L. monocytogenes* | FSL B8-0270 / PS00959 | Dairy processing facility, shoe change location | CU-11,534 | I | (Beno et al., 2016) |
| *L. monocytogenes* | FSL B8-0156 / PS00960 | Dairy processing facility, drain in adjacent room to small make room | CU-258,67 | I | (Beno et al., 2016) |
| *L. monocytogenes* | FLS B8-0053 / PS00961 | Dairy processing facility, swab, floor/wall junction in processing room | CU-29,361 | I | (Beno et al., 2016) |
| *L. monocytogenes* | FSL A5-0161 / PS00962 | Dairy processing facility, swab, underneath breakroom table | CU-54,570 | II | (Beno et al., 2016) |
| *L. monocytogenes* | FSL B8-0090 / PS00963 | Dairy processing facility, red hose in the milk receiving room | CU-133,39 | I | (Beno et al., 2016) |
| *L. monocytogenes* | FSL B8-0154 / PS00964 | Dairy processing facility, east end of south drain | CU-490,155 | II | (Beno et al., 2016) |
| *Enterococcus faecium* | 152 / PS01156 | Floor-drain of food processing plant | NA | NA | (Zhao et al., 2004) |
| *Enterococcus lactis* | C-1-92 / PS01155 | Floor-drain of food processing plant | NA | NA | (Zhao et al., 2004) |
| *Pseudomonas* spp. | PS02313 | Ice cream processing facility B | NA | NA | This study |

**FIG 1.**
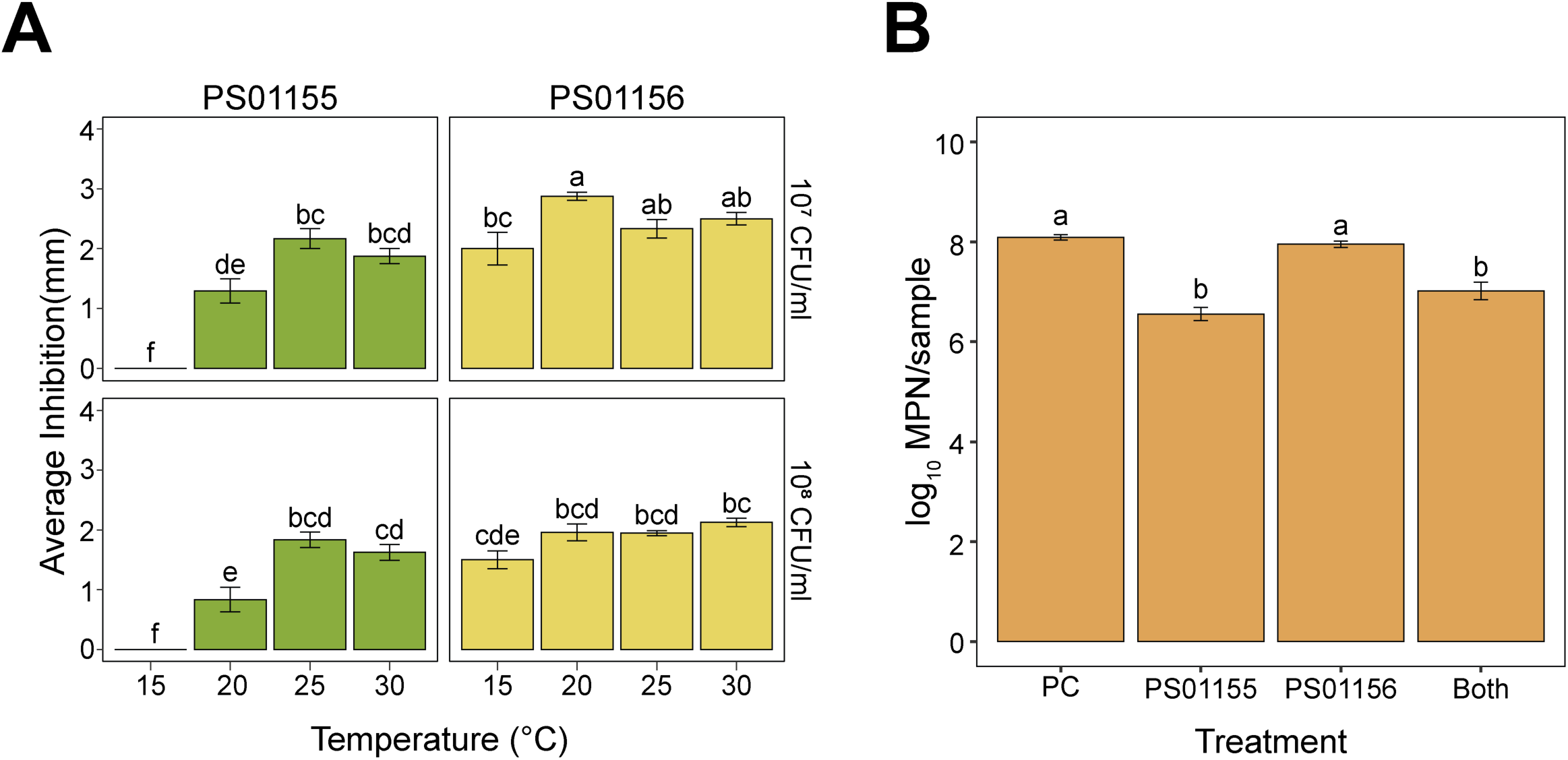
Inhibition of *L. monocytogenes* by two lactic acid bacteria strains. (A) Inhibition of *L. monocytogenes* by lactic acid bacteria strains PS01155 and PS01156 using the spot-inoculation assay. *L. monocytogenes* was spread onto lawns in two concentrations (∼10^7^ and ∼10^8^ CFU/ml, top and bottom panels, respectively) at 4 temperatures (15, 20, 25, or 30°C). (B) Inhibition of an 8-strain *L. monocytogenes* cocktail by lactic acid bacteria strains PS01155 or PS01156 or both strains together in Brain Heart Infusion broth at 15°C. A positive control (PC) tube contained only *L. monocytogenes* cocktail. Letters on top of the bars indicate significant differences (p<0.05).

The use of lactic acid bacteria as biological controls in dairy processing facilities would involve an application of cultures in food processing areas with temperatures ranging by 10-20°C. Hence, we further tested the antilisterial abilities of PS01155 and PS01156 against an 8-strain cocktail of *L. monocytogenes* strains by culturing them in polypropylene tubes containing BHI broth at 15°C for 3 days. A positive control contained only the *L. monocytogenes* cocktail to determine the reductions in attached *L. monocytogenes* concentration due to the addition of lactic acid bacteria cultures. The concentration of *L. monocytogenes* in the attached biomass after incubation was 8.09±0.05, 6.55±0.13, 7.95±0.06, or 7.01±0.18 log_10_ MPN/sample for the positive control, PS01155, PS01156, or both PS01155 and PS01156 treatments, respectively (Fig. 1B). The presence of PS01155 or both lactic acid bacteria strains together significantly reduced the concentration of *L. monocytogenes* in the attached biomass compared to the positive control by 1.54±0.14 and 1.07±0.13 log_10_ MPN/sample, respectively (Fig. 1B) (p = 1.68 × 10^−6^). However, PS01156 did not significantly reduce the concentration of *L. monocytogenes* compared to the positive control (Fig. 1B).

### The bacterial microbiota from three studied ice cream processing environments differed in taxonomic compositions

Six environmental samples were collected from the ice cream processing area of three small- to medium-scale dairy processing facilities (A, B, and C) and combined into a composite microbiome sample for each facility. To determine the baseline composition of the bacterial environmental microbiota of the composite samples collected from the studied ice cream processing facilities, we conducted amplification and sequencing of the V4 region of the 16S rRNA gene. A total of 194, 342, and 269 Amplicon Sequence Variants (ASVs) were obtained for composite samples in facilities A, B, and C, respectively. To visualize the taxonomic composition of the bacterial microbiota, present in the three ice cream processing facilities’ environments at the time of sampling, we plotted the relative abundance of the bacterial ASVs present in a relative abundance above 2% (Fig. 2). The three facilities differed in taxonomic composition of bacterial microbiota present in ice cream processing environments at the time of sampling. Facility A and B had a high relative abundance of *Pseudomonas* (26.5% in A and 48.7% in B), while in Facility C, *Pseudomonas* ASVs were detected at relative abundance below 2%. Members of *Enterobacteriaceae* were also present at high relative abundance in Facility A (8.7% *Citrobacter*) and Facility B (4.0% *Klebsiella*). In contrast, the microbiota of Facility C was composed of *Paracoccus* (25.8%), *Kocuria* (15.6%), *Corynebacterium* (7.1%), *Amaricoccus* (3.2%), *Rhodococcus* (2.4%), and *Thermomonas* (2.1%) (Fig. 2).

**FIG 2.**
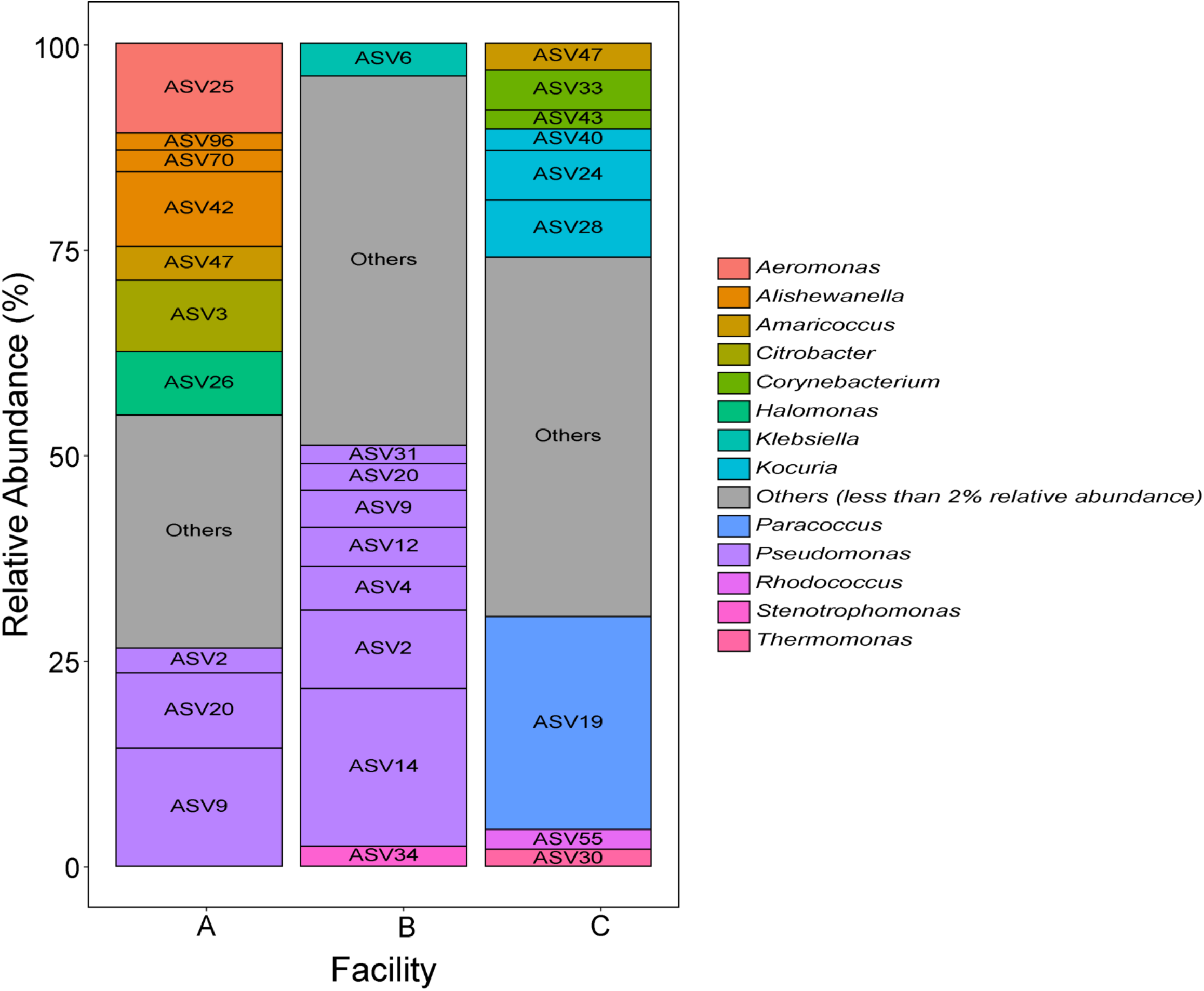
Bacterial microbiota of composite samples collected from three ice cream processing facilities. Bar plots represent the bacterial microbiota composition of the composite samples collected from three ice cream processing facilities (A, B, and C). All amplicon sequence variants (ASVs) that were present in a relative abundance above 2% in a facility are included in the plots and color coded by taxonomic genera. All taxa below the cutoff were collapsed under the “Others” category.

### *Enterococcus faecium* PS01155 and *Enterococcus lactis* PS01156 did not significantly reduce the concentration of attached *L. monocytogenes* when co-cultured with environmental microbiomes of ice cream processing facilities

Lactic acid bacteria used as biological controls need to compete for nutrients and space with the resident microbiota of food processing facilities. To determine whether *Enterococcus faecium* PS01155 and *Enterococcus lactis* PS01156 can inhibit *L. monocytogenes* in the presence of environmental microbiota of ice cream processing facilities, we conducted a competitive exclusion assay. An 8-strain cocktail of *L. monocytogenes* was co-cultured with PS01155, PS01156, or both PS01155 and PS01156 in the presence of a composite environmental microbiome collected from each ice cream processing environment in polypropylene tubes at 15°C for 3 days. A positive control (PC) contained the environmental microbiome and the *L. monocytogenes* cocktail, and a negative control (NC) contained only the environmental microbiome from each facility. The initial level of aerobic mesophilic bacteria in the environmental microbiome added to the tubes was 7.38±0.28, 5.37±0.09, and 5.38±0.11 log_10_ CFU/ml for facilities A, B, and C, respectively, and was not significantly different between facilities (Fig. 3A). After 3 days of incubation at 15°C there was no significant difference in the concentration of aerobic bacteria in the attached biomass of treatment samples for any of the three facilities when compared to the initial microbiome (Fig. 3A). The addition of PS01155 or both strains together significantly increased the concentration of aerobic mesophilic bacteria in the attached biomass when co-cultured with the microbiome of facilities A and B, compared to the negative control (p = 2.83 × 10^−7^). Compared to the positive control, when co-cultured with the environmental microbiome of facility A, the attached *L. monocytogenes* increased by 0.63±0.97, 0.51±0.68, and 0.02±0.67 log_10_ MPN/sample due to the addition of PS01155, PS01156, or both PS01155 and PS01156, respectively (Fig. 3B). When co-cultured with the environmental microbiome of facility B, the attached *L. monocytogenes* was reduced by 1.12±1.16, 1.03±1.14, and 0.83±1.09 log_10_ MPN/sample due to the addition of PS01155, PS01156, or both PS01155 and PS01156, respectively, compared to the positive control (Fig. 3B). When co-cultured with the environmental microbiome of facility C, the attached *L. monocytogenes* was reduced by 2.33±0.64, 2.80±0.79, and 2.49±1.09 log MPN/sample with the addition of PS01155, PS01156, or both PS01155 and PS01156, respectively, compared to the positive control (Fig. 3B). Nonetheless, none of these reductions were statistically significant when compared with the positive control (Fig 3B).

**FIG 3.**
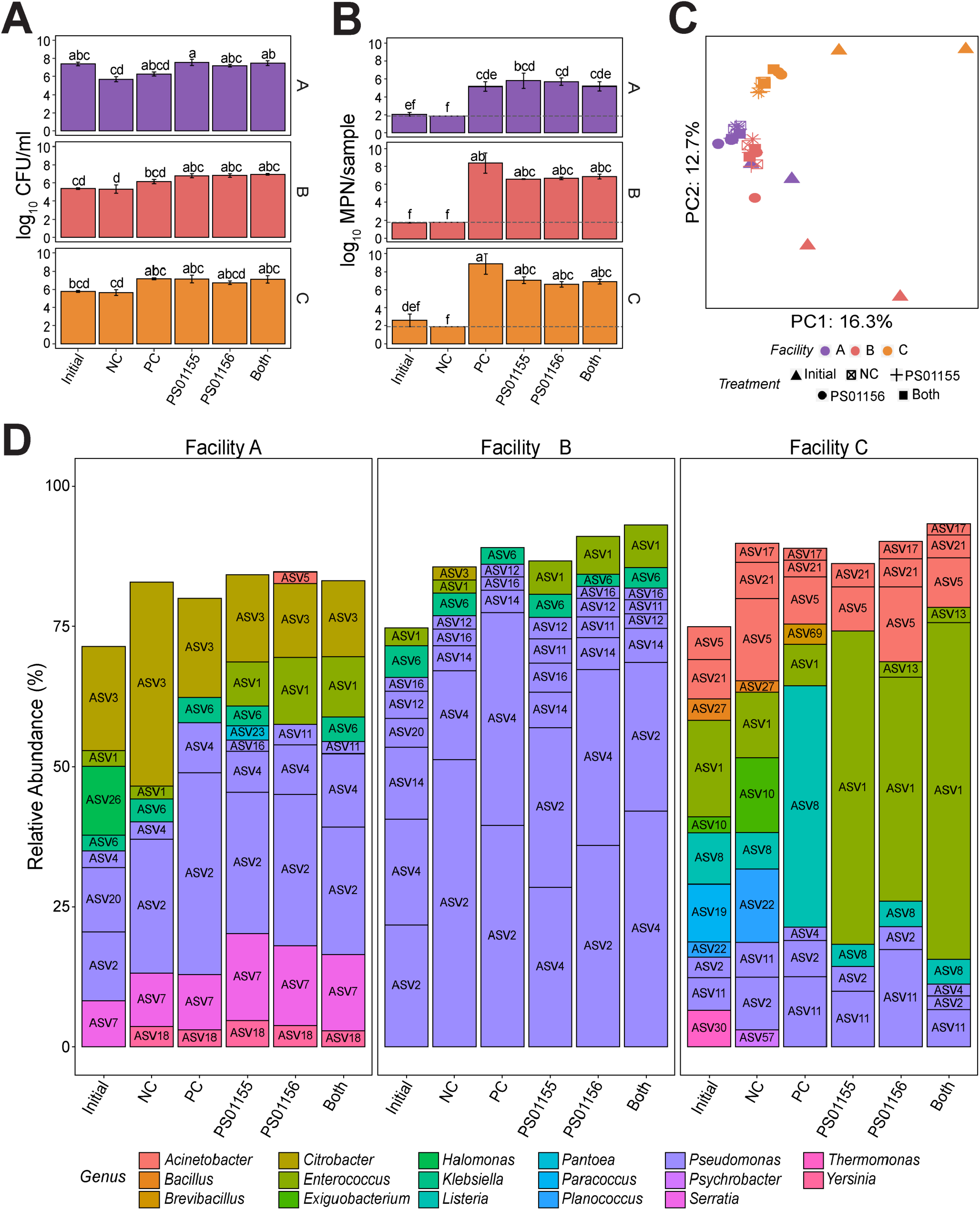
Inhibition of *L. monocytogenes* by lactic acid bacteria strains in the presence of environmental microbiota of three ice cream processing facilities. Aerobic mesophilic bacteria count (A) and *L. monocytogenes* concentration (B) of biomass from treatment samples containing lactic acid bacteria strains PS01155, PS01156, or both strains, and an 8-strain cocktail of *L. monocytogenes* attached to a polypropylene conical tube after 3 days of incubation at 15°C in the presence of the environmental microbiomes of three ice cream processing facilities (A, B, C). Bars are color coded by facility and letters above the bars represent statistical significance (p< 0.05). The limit of detection of the MPN method is shown with a grey dashed line (1.85 log_10_ MPN/sample). (C) Principal component plot of microbiota composition. Symbols indicate sample treatment and colors indicates the facility of microbiome origin. (D) Microbiota composition of attached biomass after 3 days of incubation at 15°C. Bars represent the relative abundance of amplicon sequence variants (ASVs) present in a relative abundance above 2% and are color coded by genus. Treatment labels indicate “Initial” microbiome added at the beginning of the experiment; “NC”: environmental microbiota attached biomass after 3 days at 15°C; “PC”: *L. monocytogenes* cocktail with environmental microbiota attached biomass after 3 days at 15°C; “PS01155”: *L. monocytogenes* cocktail+ PS01155+ environmental microbiota attached biomass after 3 days at 15°C; “PS01156”: *L. monocytogenes* cocktail+ PS01156 + environmental microbiota attached biomass after 3 days at 15°C, “Both”: *L. monocytogenes* cocktail+ PS01155 + PS01156 + environmental microbiota attached biomass after 3 days at 15°C.

To characterize the bacterial composition of the attached bacterial biomass of each treatment sample, we used 16S rRNA V4 amplicon sequencing. Microbial community standards were included at the DNA extraction and at the PCR amplification steps to identify potential biases introduced during these steps. All organisms included in the kit were detected after sequencing, and their relative abundances were similar to the microbial composition supplied by the manufacturer, with a few exceptions (Table S1). *Lactobacillus fermentum* had a lower relative abundance than expected, both at the DNA extraction and amplification step; while *Staphylococcus aureus* and *Bacillus subtillis* had a higher relative abundance at the DNA extraction step, suggesting a bias by the DNA extraction kit towards these organisms. Further, ASVs that did not match the composition of the standards were detected bioinformatically, suggesting contaminant DNA was added to the samples during DNA extraction, PCR amplification, library preparation, or sequencing steps. Nonetheless, contaminant sequences were removed from the dataset based on the results from the negative control samples included in the experimental design, before further analyses. PCA was used to visualize sample clustering based on microbiota composition. The first two principal components accounted for 29.2% of the variance in the dataset (PC1: 16.5%; PC2: 12.7%) and showed clustering of the samples by facility regardless of the treatment (Fig. 3C). A significant facility effect was also confirmed by PERMANOVA (p = 0.001). Further, the attached microbiota after 3 days of incubation at 15°C was significantly different from the microbiota added at the beginning of the experiment for all tested sample (p < 0.004). To visualize the taxonomic composition of the microbiota in the samples, we plotted the relative abundance of all taxa with relative abundance above 2% (Fig. 3D). The microbiome from facility A added to samples at the beginning of the experiment (labelled as “Initial”) contained a high relative abundance of *Pseudomonas* (26.7%; ASV2, ASV4, ASV20), *Citrobacter* (18.6%; ASV3), and *Halomonas* (12.3%; ASV26). After 3 days of incubation at 15°C, the attached microbiota was predominated by *Pseudomonas* (mean relative abundance of 36.8%; ASV2, ASV4, ASV11), followed by *Citrobacter* (19.3%; ASV3), and *Serratia* (12.6%; ASV7). In the treatment samples with PS01155, PS01156, or both strains added to microbiome from facility A, ASV1 (*Enterococcus*) occurred at a mean relative abundance of 10.2 %. Further, the relative abundance of ASV1 (*Enterococcus*) was 5.6, 9.7, and 8.5% higher in samples that were treated with PS01155, PS01156, or both strains, respectively, compared to the negative control (Fig. 3D). The microbiome from facility B added to samples at the beginning of the experiment (labelled as “Initial”) contained a high relative abundance of *Pseudomonas* (65.9 %; ASV2, ASV4, ASV12, ASV14, ASV16, ASV20), *Klebsiella* (5.7%; ASV6), and *Enterococcus* (3.1%; ASV1). After 3 days of incubation at 15°C, the attached microbiota were predominated by *Pseudomonas* (mean relative abundance 80.7%; ASV2, ASV4, ASV12, ASV14, ASV16), *Enterococcus* (5.7%; ASV1), and *Klebsiella* (3.4%; ASV6). In the samples with PS01155, PS01156, or both strains, added to the microbiome of facility B, ASV1 (*Enterococcus*) occurred at a mean relative abundance of 6.8 %. The relative abundance of ASV1 (*Enterococcus*) was 3.6, 4.4, and 5.4% higher in the samples that were treated with PS01155, PS01156, or both lactic acid bacteria strains, respectively, compared to the negative control (Fig. 3D). The microbiome from facility C added to samples at the beginning of the experiment (labelled as “Initial”) contained a higher diversity of taxa compared to the other two facilities, including *Enterococcus* (17.2%; ASV1), *Acinetobacter* (12.8%; ASV5, ASV21), and *Paracoccus* (10.3%; ASV19). After 3 days of incubation at 15°C, the attached microbiota was predominated by *Enterococcus* (mean relative abundance 36.1%; ASV1, ASV13), *Acinetobacter* (17.3%; ASV21, ASV5), and *Pseudomonas* (16.8%; ASV11, ASV2). In the treatment samples with PS01155, PS01156, or both strains, added to the microbiota of facility C, ASV1 (*Enterococcus*) occurred in the attached biomass at a mean relative abundance of 51.9%. The relative abundance of ASV1 (*Enterococcus*) was 44.2, 28.3, and 48.4% higher in the samples that were treated with PS01155, PS01156, or both strains, respectively, compared to the negative control (Fig. 3D).

Given that we observed lower inhibition of attached *L. monocytogenes* in samples co-cultured with microbiomes of facilities with a high relative abundance of *Pseudomonas* ASVs (i.e., facilities A and B), we further investigated whether a lower concentration of environmental microbiomes (i.e., lower concentration of *Pseudomonas*) would increase the antilisterial activity of the lactic acid bacteria strains in the attached biomass model. To test this hypothesis, we repeated the experiment using a 100-fold dilution of the environmental microbiome of facility B which resulted in a starting total aerobic bacteria concentration of 2.93±0.03 log_10_ CFU/ml. After incubation, we observed a 1.40±0.31, 0.68±0.20, and 0.74±0.29 log_10_ MPN/sample reduction of attached *L. monocytogenes* in treatment samples containing PS01155, PS01156, or a combination of both lactic acid bacteria strains, respectively. Compared to the results obtained when using the undiluted microbiomes of Facility B (Fig 3B), there was a non-significant increase in the ability of PS01155 to inhibit *L. monocytogenes*, and non-significant decrease in the ability of PS01156 or both strains combined to inhibit *L. monocytogenes* (p=0.989).

### The microbiome composition of ice cream processing facilities resulted in differences in the attachment of *Enterococcus* ASVs to polypropylene surfaces

Lactic acid bacteria intended for use as biological control strains during sanitizing of food processing facilities may need to attach to surfaces to inhibit *L. monocytogenes*. Given that we observed differences in the attached microbiota composition depending on the origin of the microbiota (i.e., Facility A, B, or C) (Fig. 3C), we hypothesized that the composition of environmental microbiota may affect the ability of the lactic acid bacteria strains to attach to surfaces. We first tested whether *Enterococcus faecium* PS01155 and *Enterococcus lactis* PS01156 could attach to the surface of polypropylene conical tubes used in our experiments. With this purpose, we incubated PS0155, PS01156, and both PS01155 and PS01156 in polypropylene tubes in BHI for 3 days at 15°C, followed by quantification of the attached biomass. The total aerobic mesophilic bacteria concentration in the attached biomass was 7.82±0.25, 5.43±0.18, and 7.16±0.07 log_10_ CFU/ml for treatment samples containing PS01155, PS01156, and both strains together, respectively (Fig. 4A). PS01155 or both strains together attached to polystyrene tubes at a significantly higher concentration compared to PS01156 (p=1.23*10^−5^) (Fig. 4A).

**FIG 4.**
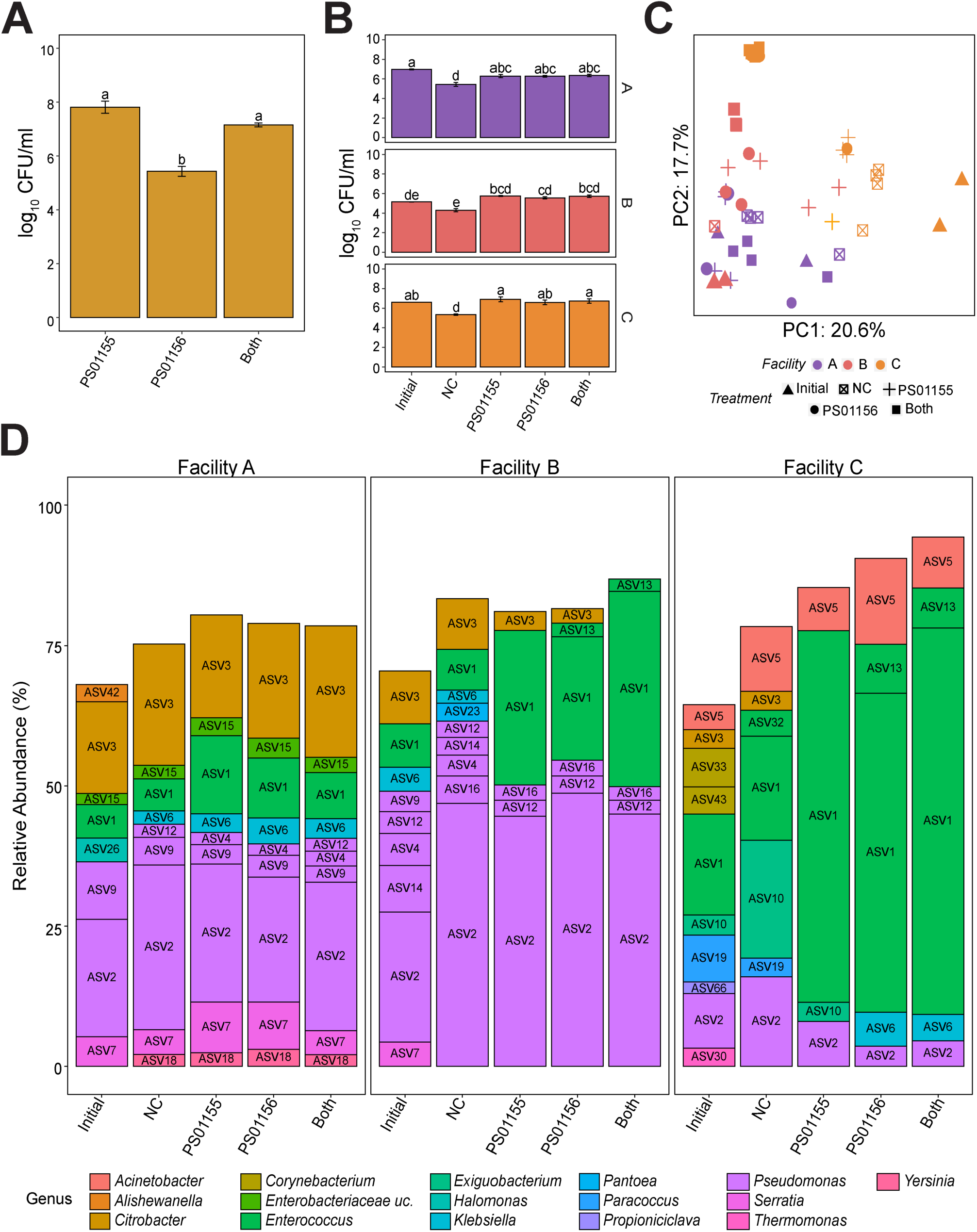
Attachment of lactic acid bacteria to polypropylene tubes in the presence of environmental microbiota of three ice cream processing facilities. (A) Attachment of lactic acid bacteria strains PS01155, PS01156, or both strains to a polypropylene tube after 3 days of incubation at 15°C. Letters above the bars represent statistical differences (p < 0.05). (B) Aerobic mesophilic count of attached microbiota containing lactic acid bacteria strains PS01155, PS01156, or both strains, attached to a polypropylene conical tube after 3 days of incubation at 15°C co-cultured with the environmental microbiota of three ice cream processing facilities (A, B, C). Bars are color coded by facility and letters above the bars represent statistical differences (p < 0.05). (C) Principal component plot of microbiota composition. Symbols indicate treatment and color indicates the facility of microbiota. (D) Microbiota composition of attached biomass after 3 days of incubation at 15°C. Bars represent the relative abundance of amplicon sequence variants (ASVs) present in a relative abundance above 2% and are color coded by genus. Treatment labels indicate “Initial” microbiota added at day zero; “NC”: environmental microbiota attached biomass after 3 days at 15°C; “PS01155”: PS01155+ environmental microbiota attached biomass after 3 days at 15°C; “PS01156”: PS01156 + environmental microbiota attached biomass after 3 days at 15°C, “Both”: PS01155 + PS01156 + environmental microbiota attached biomass after 3 days at 15°C.

We further investigated the extent to which PS01155 and PS01156 attached to polypropylene tubes in the presence of the environmental microbiomes of three ice cream processing environments. PS01155, PS01156, or both PS01155 and PS01156 were cultured in the presence of a composite environmental microbiome collected from each ice cream processing facility in polypropylene tubes at 15°C for 3 days. A negative control (NC) contained only the environmental microbiome from each facility. The initial concentration of aerobic mesophilic bacteria in the environmental microbiome added to the tubes was 6.97±0.04, 5.15±0.02, and 6.61±0.01 log_10_ CFU/ml for facilities A, B, and C, respectively (Fig. 4B). After 3 days of incubation, we did not observe a significant effect of the facility microbiome on the aerobic mesophilic bacteria in the attached biomass of any treatment (Fig. 4B). However, compared to the negative control, the addition of PS01155, PS01156, or both strains together to microbiomes from facilities A, B, or C significantly increased the concentration of aerobic mesophilic bacteria in the attached biomass (p = 1.46 × 10^−13^) (Fig. 4B).

To characterize bacterial composition of the attached biomass, we used 16S rRNA V4 gene region amplicon sequencing. PCA was used to visualize the similarity of microbiota of each treatment sample in a two-dimensional space. The first two principal components accounted for 38.3% of the variance in the dataset (PC1: 20.6%; PC2: 17.7%) and showed clustering of samples by facility, regardless of the treatment (Fig. 3C), which was confirmed by PERMANOVA (p = 0.001). To visualize taxonomic composition of the microbiota in samples, we plotted the relative abundance of all taxa present with a relative abundance above 2%(Fig. 4D). The microbiome from facility A added to samples at the beginning of the experiment (“Initial”) contained a high relative abundance of *Pseudomonas* (31.2%; ASV2, ASV9), *Citrobacter* (16.3%; ASV3), *Enterococcus* (6.0%; ASV1), *Serratia* (5.3%; ASV7), *Halomonas* (4.2%; ASV26), *Alishewanella* (3.1%; ASV42), and *Enterobacteriaceae* unclassified (2.0%; ASV15). After 3 days of incubation at 15°C, the attached microbiota continued having a high mean relative abundance of *Pseudomonas* (32.3%; ASV2, ASV4, ASV9), *Citrobacter* (21.0%; ASV3), *Enterococcus* (9.6%; ASV1), *Serratia* (6.5%; ASV7), *Enterobacteriaceae_*unclassified (3.0%; ASV15), and additionally contained over 2% of *Klebsiella* (3.4%; ASV6) and *Yersinia* (2.4%; ASV18). The relative abundances of *Halomonas* and *Alishewanella* decreased below 2%. The relative abundance of ASV1 (*Enterococcus*) increased by 8.2, 5.0, and 2.5 % in samples that were treated with PS01155, PS01156, or both strains, respectively, compared to the negative control (Fig. 4D).

The microbiome from facility B added to samples at the beginning of the experiment (“Initial”) contained a high relative abundance of *Pseudomonas* (44.7%; ASV2, ASV4, ASV12, ASV14, ASV16), *Citrobacter* (9.4%; ASV3), *Enterococcus* (7.8%; ASV1), Serratia (4.3%; ASV7), and *Klebsiella* (4.3%; ASV6). After 3 days of incubation at 15°C, the attached microbiota continued having a high mean relative abundance of *Pseudomonas* (54.1%; ASV2, ASV12, ASV16), *Enterococcus* (24.0%; ASV1), *Citrobacter* (4.9%; ASV3). The relative abundance of *Pantoea* (3.2%; ASV23) increased above 2% and that of *Klebsiella* and *Serratia* decreased below 2 %. The relative abundance of ASV1 (*Enterococcus*) increased by 20.2, 14.7, and 27.5% in the samples that were treated with PS01155, PS01156, or both strains, respectively, compared to the negative control (Fig. 4D).

The microbiome from facility C added to samples at the beginning of the experiment (“Initial”) contained a high relative abundance of the genera *Enterococcus* (18.0%; ASV1), *Corynebacterium* (11.7%; ASV 33, ASV43), *Pseudomonas* (9.7%; ASV2), *Paracoccus* (8.4%; ASV19), *Acinetobacter* (4.5%; ASV5), *Exiguobacterium* (3.6%; ASV10), *Citrobacter* (3.3%; ASV3), *Thermomonas* (3.2%; ASV30), and *Propioniciclava* (2.1%; ASV66). After 3 days of incubation at 15°C, the attached microbiota continued having a high mean relative abundance of *Enterococcus* (57.8%; ASV1), *Exiguobacterium* (12.2%; ASV10), *Acinetobacter* (10.9%; ASV5), *Pseudomonas* (8.0%; ASV2), *Citrobacter* (3.4%; ASV3), and *Paracoccus* (3.3%; ASV19). The relative abundance of *Klebsiella* (5.3%; ASV6) increased above 2%, whereas that of *Thermomonas* and *Propioniclava* decreased below 2%. The relative abundance of ASV1 (*Enterococcus*) increased by 47.7, 38.3, and 50.3% in the samples that were treated with PS01155, PS01156, or both strains, respectively, compared to the negative control (Fig. 4D).

### The presence of *Pseudomonas* decreases the antilisterial ability of *Enterococcus faecium* PS01155

We further investigated whether the presence of *Pseudomonas* decreases the antilisterial ability of *Enterococcus faecium* PS01155 and *Enterococcus lactis* PS01156. For this purpose, we selectively isolated *Pseudomonas* spp. from the environmental microbiome sample from Facility B and confirmed their identity using Sanger sequencing of the 16S rRNA gene region. We extracted the V4 regions from 16S rRNA Sanger sequences of the isolated *Pseudomonas* and compared their sequence to the ASVs that were taxonomically classified as *Pseudomonas* in our prior experiment. To test the effect of the presence of *Pseudomonas* on the antilisterial abilities of the two lactic acid bacteria, we selected *Pseudomonas* PS02313 as it had 0 SNPs with ASV2, the *Pseudomonas* ASV with the highest relative abundance in the attached microbiota of facility B after treatment with lactic acid bacteria strains (Fig 3D). An 8-strain *L. monocytogenes* cocktail was co-cultured with PS01155, PS01156, or both PS01155 and PS01156 in the presence of *Pseudomonas* (∼1.0*10^3^ CFU/ml) in polypropylene tubes at 15°C for 3 days. The concentration of *Pseudomonas* in the experiment was equal to that present in the composite microbiome collected from Facility B. A positive control (PC) contained only the *L. monocytogenes* cocktail. In the presence of *Pseudomonas* PS02313, there was a 0.18±0.14, 0.20±0.06, and 0.10±0.06 log_10_ MPN/sample reduction in the concentration of attached *L. monocytogenes* when co-cultured with PS01155, PS01156, or both strains together, respectively, compared to the positive control. Further, the presence of *Pseudomonas* PS02313 significantly reduced the antilisterial ability of PS01155 or both strains by 1.36±0.20 and 0.97±0.10 log_10_ MPN/sample, respectively, compared to the reduction of attached *L. monocytogenes* without the presence of *Pseudomonas* (Fig 5).

**FIG 5.**
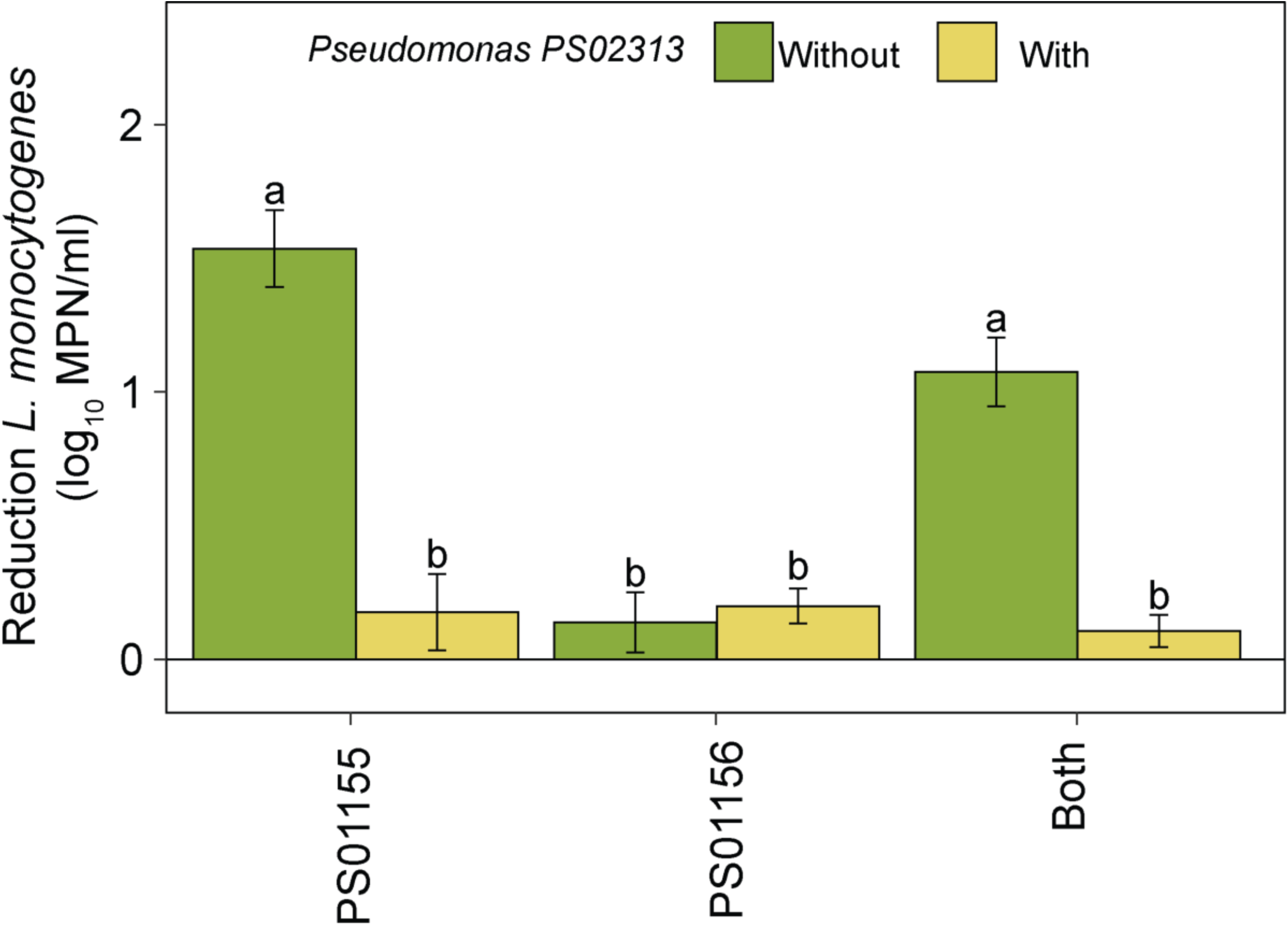
Reduction of attached *L. monocytogenes* concentration in the presence or absence of *Pseudomonas* PS02313. Reduction of attached *L. monocytogenes* concentration by lactic acid bacteria strains PS01155, PS01156, or both strains, when co-cultured in the presence or absence of *Pseudomonas* PS02313 in polypropylene tubes for 3 days at 15°C. Letters above the bars represent statistical significance (p < 0.05). Bars are color coded by the presence or absence of *Pseudomonas* PS02313 in the cultures.

## DISCUSSION

### The environmental microbiota collected from three ice cream processing facilities differed among facilities

Environmental samples were collected from three ice cream processing facilities during production times, from different locations that can be potential harborage sites for *L. monocytogenes* (e.g.: drains, cracks on floors). When comparing the hygiene of the same locations across facilities, A and B had visible soils at the time of sampling, particularly in drains and drain covers, whereas facility C appeared clean (i.e., no soils or milk was visible) during visual inspection. We did not carry out extensive surveillance of cleaning and sanitation procedures conducted within sample facilities. However, the observable buildups (i.e., food debris) in unclean spots could explain the differences observed in the composition of the microbiota among facilities.

The overall composition of the bacterial microbiota of the three ice cream processing facilities was different. Facility A and B, which appeared less clean based on visual observation, had a high relative abundance of *Pseudomonas* and *Enterobacteriaceae* (e.g., *Citrobacter, Klebsiella*). Members of the *Enterobacteriaceae* family are commonly used as indicators of cleaning and sanitizing effectiveness in food processing environments (Buchanan & Oni, 2012; Hervert et al., 2016). *Pseudomonas* are known food spoilage organisms that can be introduced to the food processing environment from soil or water (Raposo et al., 2017). Given their psychrotrophic nature and their ability to adhere, they can establish biofilms in hard-to-clean areas (Mann & Wozniak, 2012). Given the dynamic nature of microbiota in food processing facilities (Johnson et al., 2021), we do not purport to provide a thorough description of the bacteria residing within these food environments, but rather a snapshot of the microbiota collected for our experiments. Further longitudinal studies are needed to determine the spatial and temporal microbiota of ice cream processing facilities, whether there is core microbiota shared among facilities, and whether there is an association between environmental microbiota and quality or safety issues.

### Lactic acid bacteria strains PS01155 and PS01156 did not significantly inhibit *L. monocytogenes* when co-cultured with environmental microbiomes of ice cream processing facilities

Due to their inhibitory characteristics against *L. monocytogenes*, lactic acid bacteria may be used to complement cleaning and sanitizing operations within dairy processing facilities as biological control agents (Camargo et al., 2018). Two lactic acid bacteria strains selected for this study (i.e., PS01155 and PS01156) were whole genome sequenced and identified as *Enterococcus faecium* and *Enterococcus* lactis, respectively (Sinclair et al., 2022). *Enterococcus* species may inhibit *L. monocytogenes* by producing bacteriocins, sometimes referred to as enterocins. Enterocins are low molecular weight, ribosomally synthesized, heat-stable peptides, that form pores in the cell membrane of susceptible organisms (ben Braïek & Smaoui, 2019). The production of bacteriocins by *Enterococcus* spp. is affected by temperature, *L. monocytogenes* concentration, growth media, and growth stage of the producing strain (Schirru et al., 2014). In this study we observed a temperature-dependent inhibition of *L. monocytogenes* by strains PS01155 and PS01156, with both strains showing less inhibition at lower temperatures, using the spot-inoculation method. Schirru et al., (2014) showed that the inhibition of *L. monocytogenes* by *Enterococcus faecium* at different temperatures was dependent on the strain, with two strains of *E. faecium* showing an increased inhibition at higher incubation temperatures, while other strains did not show this trend (Schirru et al., 2014). In contrast to our results, Sinclair *et al*., (2022) showed an increase in the inhibitory activity of PS01155 against *L*.*monocytogenes* strains isolated from tree fruit packing facilities at lower temperatures (i.e., 15 and 20°C).

In this study, we did not observe a significant reduction of *L. monocytogenes* when co-cultured with lactic acid bacteria strains PS01155 or PS01156 and environmental microbiomes of ice cream processing facilities. However, the concentration of *L. monocytogenes* used in the assays was high (∼3*10^6^ CFU/ml CFU/ml) and represented a worst-case scenario. Given that the concentration of *L. monocytogenes* may affect the antilisterial performance of lactic acid bacteria (Sinclair et al., 2022), it is likely that these high concentrations of *L. monocytogenes* may have hindered the antilisterial activities of the lactic acid bacteria. In contrast to our results, Zhao *et al*. (2006) observed a ∼2.5 log reduction of *Listeria* spp. when both lactic acid bacteria strains were used on drains in rooms at ∼15°C within a poultry processing facility (Zhao et al., 2006). Differences between the two studies could be due to the native microbiota of poultry processing facilities or to differences in the susceptibility of different *Listeria* spp. strains to the antilisterial effect of lactic acid bacteria. Further, our results are limited to the model system applied, thus further work needs to be conducted to assess the antilisterial effectiveness of lactic acid bacteria within a real-world scenario, on dairy processing equipment-relevant materials (e.g., stainless steel, concrete, rubber), and a lower concentration of *L. monocytogenes*.

### The inhibition of *L. monocytogenes* by lactic acid bacteria strains PS01155 and PS01156 was influenced by the environmental microbiota of ice cream processing facilities

In this study, we observed differences in the reduction of *L. monocytogenes* attached to a polypropylene surface depending on the composition of the accompanying microbiota. Further, a greater inhibition of *L. monocytogenes* was observed when strains PS01155 and PS01156 were co-cultured with the environmental microbiome sample obtained from facility C, compared to A and B. The attached microbiota of samples co-cultured with lactic acid bacteria and the environmental microbiome sample from facility C contained a lower relative abundance of *Pseudomonas* and *Enterobacteriaceae*, compared that of facilities A and B. *Pseudomonas* are psychrotrophs that can attach to surfaces to form biofilms (Mann & Wozniak, 2012). Thus, it is likely that in samples containing microbiomes from facilities A and B, *Pseudomonas* attached to the polypropylene surface faster than the lactic acid bacteria, decreasing the concentration of PS01155 and PS01156 on the surface, which resulted in reduced antilisterial activity. Furthermore, species of *Enterobacteriaceae*, including *Citrobacter* and *Klebsiella*, can form biofilms within ice cream processing environments, suggesting that a similar mechanism might be occurring. It is also likely that some members of the microbiota of facility C may promote attachment of the lactic acid bacteria by an unknown mechanism. We further determined that the presence of one *Pseudomonas* spp. decreased the antilisterial abilities of PS01155. We hypothesize that the presence of psychrotrophic biofilm formers may prevent the attachment of lactic acid bacteria to surfaces through competitive exclusion. Nonetheless, other mechanisms of protection of *L. monocytogenes* are possible. For example, previous research has observed that *L. monocytogenes* can promote biofilm formation by *Pseudomonas fluorescens* (Puga et al., 2018). Additional research is needed to understand whether other *Pseudomonas* spp., and other microbiota such as members of the *Enterobacteriaceae* family, affect the antilisterial abilities of lactic acid bacteria. Further, in this study, we did not determine the fungal microbiota present in microbiomes of ice cream processing facilities and their effect on the inhibition of *L. monocytogenes*. To the best of our knowledge, there have been no studies that investigated the attachment of lactic acid bacteria to abiotic surfaces when in the presence of resident bacteria of food processing environments. Further work is therefore needed to evaluate the kinetics of attachment of PS01155 and PS01156 when in the presence of certain environmental microbiota. Understanding the mechanisms by which environmental microbiota may decrease the efficacy of biological controls will promote an effective use of lactic acid bacteria within cleaning and sanitizing operations.

### Conclusions

Lactic acid bacteria proposed as biological control agents in food processing environments need to compete for space and nutrients with the resident microbiota present in food processing facilities, which may affect their antilisterial activity. Our study showed that the composition of the microbiota collected from ice cream processing facilities and the presence of certain microbial taxa affect the attachment and inhibitory effect of lactic acid bacteria strains against *L. monocytogenes*. Further studies are needed to evaluate whether certain taxa effectively inhibit or reduce the antilisterial properties of lactic acid bacteria, and the mechanisms behind microbiota interactions.

## MATERIALS AND METHODS

### Bacterial strains

Two lactic acid bacteria strains, *Enterococcus faecium* PS01155 and *Enterococcus lactis* PS01156, were purchased from the American Type Culture Collection (ATCC, Manassas, VA) (Table 1). The two lactic acid bacteria strains used in this study were selected due to their reported inhibition of *L. monocytogenes* in poultry processing facilities, where a ∼2.5 log CFU/100 cm^2^ reduction of *Listeria* spp. was observed in drains located in rooms with an ambient temperature of ∼15°C (Zhao et al., 2006). *L. monocytogenes* strains previously isolated from small-scale dairy processing facilities (Beno et al., 2016) were obtained from Cornell University. These isolates were selected to represent a diverse set of *L. monocytogenes* genotypes representing two lineages with different PGFE patterns (Table 1). All bacterial isolates were cryopreserved in Brain Heart Infusion (BHI) broth (BD, cat. no. 237500, Franklin Lakes, NJ) supplemented with 20% v/v sterile glycerol at -80°C.

### Bacterial culture preparation

All bacteria strains were streaked from cryostocks onto BHI agar and incubated at 35°C for 24 hours. Colonies from each strain were suspended in 9 ml of sterile 0.1% peptone water (BD, cat. no. 218071, Franklin Lakes, NJ) and adjusted to an optical density at 600 nm of 0.13 (∼10^8^ CFU/ml), prior to use as described in subsequent sections.

### Assessment of antilisterial activity of lactic acid bacteria strains

Spot-inoculation assays (Kim et al., 2016) were performed to determine whether the two lactic acid bacteria were inhibitory to *L. monocytogenes* strains (N = 8) previously isolated from dairy processing environments at 4 different temperatures. Lawns of each *L. monocytogenes* strain were prepared by streaking ∼100 µl of cultures (prepared in concentrations ∼10^7^CFU/ml and ∼10^8^CFU/ml) onto BHI agar plates. One microliter (∼10^5^ CFUs) of each lactic acid bacteria strain culture (PS01155 and PS01156) was spot inoculated onto the lawn of *L. monocytogenes* in triplicates and incubated at 15, 20, 25, or 30°C for 3 days. After incubation, the zones of inhibition produced by each spot of lactic acid bacteria strains were measured from the border of the colony-spot to the outer edge of the inhibition zone at three locations using a ruler. The experiment was conducted in three independent replicates. One-way Analysis of Variance (ANOVA) was used to assess the effect of temperature on the antilisterial abilities of PS01155 and PS01156 using the R package stats v4.1.0 (R Core Team, 2021), followed by Tukey’s Honest Significant Differences test using the R package agricolae (de Mediburu, Felipe, 2021) in R v4.1.0 (R Core Team, 2021).

To determine whether the lactic acid bacteria strains could inhibit *L. monocytogenes* when grown in broth and attached to an abiotic surface, an 8-strian cocktail of *L. monocytogenes* (∼3*10^6^ CFU/ml) was co-cultured with PS01155, PS01156 or both PS01155 and PS01156 (∼6*10^7^ CFU/ml) in polypropylene conical tubes (cat. no. 89039-670, VWR, Radnor, PA) in BHI broth. In addition, a positive control (PC) tube containing only the *L. monocytogenes* cocktail and a negative control (NC) tube containing just BHI broth were included in the experiment. All tubes were incubated statically at 15°C for 3 days. The concentration of *L. monocytogenes* was determined in the attached biomass as described in a subsequent section. The experiment was conducted in two independent biological replicates, each with two technical replicates per treatment. One-way ANOVA was used to assess statistical differences in the attachment of PS01155 and PS01156, followed by Tukey’s Honest Significant Differences test, as previously described.

### Environmental microbiome collection and preservation

Three small- to medium-scale dairy processing facilities in the Northeastern U.S. were selected to participate in the study. Each facility was visited once during Fall 2019 and six environmental samples were collected in each facility from non-food contact areas inside the ice cream manufacturing room, during processing hours. Four of the samples were collected from similar areas in the ice cream processing room of each facility, including drains adjacent to the mix holding tank and the ice cream freezer, and floors by the mix holding tank and by the ice cream freezer. Two additional samples were collected within each facility from potential high-risk locations for *L. monocytogenes* harborage, including wheels of a cart used in the processing room and a squeegee in facility A, a floor with a visible crack and a floor mat in facility B, and the drains by the pasteurized mix tank and floor underneath the packaging area in facility C. Each sample was collected by swabbing a 40 × 40 cm area (or an equivalent area for irregular surfaces) with a 3M Hydrated Sponge with Neutralizing buffer (3M, Catalog#: HS10NB2G, Saint Paul, MN). Samples were stored in a cooler, transported to the laboratory, and processed within 24 hours of collection. At the time of sampling, the temperature in the ice cream processing areas was recorded to determine the incubation temperature used in the subsequent assays detailed below.

Upon arrival to the laboratory, 90 ml of sterile BHI broth were added to each sample bag and each bag was stomached at 230 rpm for 7 minutes to release cells from a sponge. Following homogenization, the broth from the six samples collected at the same facility was combined into a sterile bottle and mixed for 5 minutes to create a composite microbiome sample for each facility. To characterize the bacterial composition of the composite microbiome sample, 50 ml of each facility’s sample were transferred to a sterile conical tube, centrifuged at 11,000 g for 20 minutes at 4°C to precipitate the cells and soils, followed by removal of the supernatant and storage of the precipitate at -80°C until DNA extraction using Omega Soil DNA kit (Omega BioTek, cat. no. D5625-01, Norcross GA) following the manufacturer’s protocol. Composite microbiome samples from each facility were cryopreserved by adding 20% v/v sterile glycerol in 50 ml conical tubes at -80°C until further use. Before the start of each experiment, composite microbiome samples were thawed at room temperature.

### Assessment of antilisterial activity of two lactic acid bacteria strains in the presence of environmental microbiomes of ice cream processing facilities

To determine whether the two lactic acid bacteria strains could inhibit *L. monocytogenes* in the presence of environmental microbiomes of ice cream processing facilities, an 8-strain cocktail of *L. monocytogenes* (∼3*10^6^ CFU/ml) was co-cultured with PS01155, PS01156 or both PS01155 and PS01156 (∼6*10^7^ CFU/ml) in polypropylene conical tubes in BHI broth, with the addition of the composite microbiome sample collected from each ice cream processing facilities. A negative control (NC) tube contained only the microbiomes from each facility, and a positive control (PC) tube contained the microbiome from each facility and the *L. monocytogenes* cocktail. All tubes were incubated statically at 15°C for 3 days. After incubation, the total aerobic mesophilic bacteria and *L. monocytogenes* concentrations were quantified in the attached biomass (Sinclair et al., 2022) as described in the subsequent section “Total aerobic mesophilic bacteria and *L. monocytogenes* quantification in attached biomass”. One-way ANOVA was used to assess statistical differences in total aerobic mesophilic bacteria and *L. monocytogenes* concentrations by lactic acid bacteria treatments, followed by Tukey’s Honest Significant Differences test. To characterize the microbiota composition of the attached biomass of each treatment, 1 ml of attached biomass was transferred to a microcentrifuge tube and frozen until DNA extraction, PCR amplification, and sequencing, as described in the subsequent section “DNA extraction, amplicon sequencing, and bioinformatic analyses”. The experiment was conducted in two independent biological replicates for each facility, each with two technical replicates per treatment.

### Attachment of lactic acid bacteria strains to polypropylene tubes in the presence of environmental microbiome from ice cream processing facilities

To determine whether the two lactic acid bacteria strains could effectively attach to a polypropylene surface, PS01155, PS01156, or both PS01155 and PS01156 (∼6*10^7^ CFU/ml) were incubated in polypropylene conical tubes containing BHI broth at 15°C for 3 days. After incubation, the total aerobic mesophilic bacteria concentration was determined in the attached biomass as described in the subsequent section “Total aerobic mesophilic bacteria and *L. monocytogenes* quantification in attached biomass”. One-way ANOVA was used to assess statistical differences in attachment of PS01155 and PS01156, followed by Tukey’s Honest Significant Differences test. To further determine whether the lactic acid bacteria could attach to the polypropylene surface in the presence of the resident microbiota of ice cream processing facilities, PS01155, PS01156, or both PS01155 and PS01156 (∼6*10^7^ CFU/ml) were co-cultured in polypropylene conical tubes in BHI broth with environmental microbiome collected from individual ice cream processing facilities. A negative control (NC) tube containing only the microbiome from each individual facility was included in the experiment. All tubes were incubated statically at 15°C for 3 days. After incubation, total aerobic mesophilic bacteria concentration was determined in the attached biomass as described in subsequent sections. One-way ANOVA was used to assess statistical differences in the attachment of PS01155 and PS01156, followed by Tukey’s Honest Significant Differences test. To characterize the bacterial microbiota composition of each treatment, 1 ml of attached biomass was transferred to a microcentrifuge tube and was kept frozen until DNA extraction, PCR amplification, and sequencing, as described in the subsequent section “DNA extraction, amplicon sequencing, and bioinformatic analyses”. The experiment was conducted in two independent biological replicates for each facility, each with two technical replicates per treatment.

### Assessment of antilisterial activity of two lactic acid bacteria strains in the presence of *Pseudomonas* spp. isolated from ice cream processing facilities

*Pseudomonas* spp. were enumerated and isolated from a composite microbiome sample collected from Facility B, by serial dilution in 0.1% peptone water and plating onto *Pseudomonas* agar (Oxoid, cat. no. CM0559B, Hampshire, U.K.) with the addition of Cetrimide-Fucidin-Cephalosporin (C-F-C) selective supplement (Oxoid, cat. no. SR0103E, Hampshire, U.K.) and incubated at 30°C for 48 hours. Sixteen colonies of different morphologies were picked, sub-streaked onto BHI agar for purification, and cryo-preserved in BHI broth supplemented with 20% v/v sterile glycerol. To identify each putative *Pseudomonas* isolate, we PCR-amplified the 16S rRNA gene region using primers 27F [AGAGTTTGATCMTGGCTCAG] (Integrated DNA Technologies, Newark, NJ) and 1495R [GGTTACCTTGTTACGACTT] (Integrated DNA Technologies, Newark, NJ), as previously described by Pomastowski *et al*., (2019). PCR amplicons were cleaned up using Exonuclease I and Shrimp Alkaline Phosphatase (Applied Biosystems, cat. no. 78201.1, Waltham, MA) and incubating at 37°C for 45 minutes, then heating to 80°C for 15 minutes, followed by cooling to 4°C. Clean PCR amplicons were sent to Penn State Genomics Core Facility (University Park, PA) for Sanger sequencing with forward primer 27F. Sequences were aligned to BLAST nucleotide database (Altschul et al., 1990) to verify the taxonomic identity of the isolates. Muscle was used in MEGA11 (Tamura et al., 2021) to align and calculate the number of base differences between Sanger sequences of *Pseudomonas* isolates and sequences of ASVs assigned to the genus *Pseudomonas*.

To determine the antilisterial ability of two lactic acid bacteria strains in the presence of *Pseudomonas* spp. isolated from ice cream processing facility B, an 8-strain cocktail of *L. monocytogenes* (∼3*10^6^ CFU/ml) was co-cultured with PS01155, PS01156, or both PS01155 and PS01156 (∼6*10^7^ CFU/ml) in polypropylene conical tubes in BHI broth, with the addition of *Pseudomonas* PS02313 (∼1.0*10^3^ CFU/ml). The positive control (PC) tube contained only the *L. monocytogenes* cocktail. All tubes were incubated statically at 15°C for 3 days. After incubation, *L. monocytogenes* concentrations were quantified in the attached biomass (Sinclair et al., 2022) as described in the subsequent section “Total aerobic mesophilic bacteria and *L. monocytogenes* quantification in attached biomass”. The experiment was conducted in two independent biological replicates, each with two technical replicates per treatment. One-way ANOVA was used to assess statistical differences in reductions of *L. monocytogenes* concentration by lactic acid bacteria treatments, followed by Tukey’s Honest Significant Differences test.

### Total aerobic mesophilic bacteria and *L. monocytogenes* quantification in attached biomass

After incubation of the conical tubes, the media containing non-attached cells was removed and loosely attached cells were removed by gently rinsing with 0.1% peptone water twice (Sinclair et al., 2022). The attached cells were resuspended in 0.1% peptone water and 2 g of 3 mm glass beads (Corning Life Sciences, cat. no. 7268-3, Corning, NY) were added to each tube, followed by 30 seconds of vortexing at a maximum speed to release the attached biomass. Total aerobic mesophilic bacteria were quantified for the initial environmental microbiota used in each experiment (labeled as “Initial”) and for the attached biomass of each treatment. The concentration of aerobic mesophilic bacteria was determined by serially ten-fold diluting a sample in 0.1% peptone water and spread plating dilutions onto BHI agar, in duplicate, followed by incubation at 35°C for 3 days. To quantify *L. monocytogenes* concentration in the attached biomass, a modified FDA Bacteriological Analytical Manual (BAM) Most Probable Number (MPN) (Hitchins, Anthony D. et al., 2022) method was used. Briefly, each treatment sample and the initial environmental microbiome suspension were serially ten-fold diluted seven times in 0.1% peptone water. Then, 100 μl of each dilution were inoculated into 900 μl of Buffered *Listeria* Enrichment broth (BLEB) (cat. no. C7803, Criterion, Hardy Diagnostics, Santa Maria, CA), in triplicates, followed by incubation at 30°C for 4 hours. After 4-hour incubation, 4 μl of selective *Listeria* enrichment supplement (90 mg acriflavine, 450 mg cycloheximide, and 360 mg sodium nalidixic acid in 40 ml of sterile water) (Sigma Aldrich, Saint Louis, MO) were added to each tube, followed by incubation at 30°C for another 44 hours. After incubation, a loopful of each sample was streaked onto Agar *Listeria* Ottavani & Agosti (ALOA) and plates (BioMerieux, Marcy-l’Etoile, France) and incubated at 37°C for 48 hours. The presence of blue colonies with a halo on ALOA plates indicated the presence of *L. monocytogenes*. The number of MPN tubes positive for *L. monocytogenes* was recorded, and the final concentration was calculated using the MPN calculator downloaded from the FDA BAM Appendix 2 (Blodgett, 2020).

### DNA extraction, amplicon sequencing, and bioinformatic analyses

Frozen attached biomass samples were thawed at room temperature and centrifuged at 11,000 g for 20 minutes to precipitate the cells. The pellet was resuspended in the lysis buffer provided with the DNeasy Biofilm kit (Qiagen, cat. no. 24000-50, Hilden, Germany) and transferred to the lysis tube, followed by DNA extraction according to the manufacturer’s protocol. Two controls were included at the DNA extraction step: a negative control to account for microbial contaminants present in DNA extraction kits or introduced during the extraction (Eisenhofer et al., 2019) and a microbial community standard (Zymo Research, cat. no. D6300, Irving, CA). The DNA concentration was quantified with Qubit 3 using the high sensitivity dsDNA kit (Invitrogen, cat. no. Q33239, Waltham, MA) fluorometer. Extracted DNA was sent to Novogene Co. (Beijing, China) for PCR amplification of the V4 region of the 16S rRNA gene and sequencing using an Illumina NovaSeq (Illumina, San Diego, CA) in two sequencing runs. A microbial community DNA standard was included with each sequencing batch (Zymo Research, cat. no. D6305, Irving, CA) to identify biases at the PCR amplification, library preparation and sequencing steps.

Sequence reads were analyzed with the R package DADA2 v3.14 pipeline by following the standard protocol for 16S rRNA V4 region amplicon sequence reads (Callahan et al., 2016). Briefly, low-quality sequence reads were filtered out, low-quality bases were trimmed, error rates were calculated for the dataset, and amplicon sequence variants (ASVs) were inferred from the remaining sequence reads. Paired-end sequence variants were merged into contigs and contigs shorter than 251 bp or longer than 253 bp were discarded. Chimeras were detected and removed, and remaining ASVs were assigned taxonomy using the Silva database v132 (Quast et al., 2013). ASVs assigned to Mitochondria or Chloroplasts were removed from the dataset before further analyses. The R package decontam v1.12.0 (Davis et al., 2018) was used to detect contaminant reads based on the negative control, using the prevalence mode with a 0.5 threshold level. A compositional analysis approach was used to analyze the microbiota data (Gloor et al., 2016, 2017). Briefly, all ASVs with zero count value in a sample were replaced with a small, non-zero value using the R package zCompositions v1.3.4 (Palarea-Albaladejo & Martín-Fernández, 2015), followed by center log ratio (CLR) transformation. Principal Component Analysis (PCA) was performed on log-transformed data to visualize microbiota composition by treatment and facility. Aitchison distances were calculated from CLR-transformed data and Permutational Multivariate Analysis of Variance (PERMANOVA) was conducted to assess statistical significance of the effect of the facility and treatment using the R packaga pairwiseAdonis v0.01 (Martinez Arbizu, 2017). To visualize the taxonomic composition of the microbiota, ASVs that had a relative abundance above 2% were plotted using the R package ggplot2 v3.3.5 (Wickham, Hadley, 2016). All analyses were carried out in R v4.1.0 (R Core Team, 2021).

## Declarations

### Ethics approval and consent to participate

Not applicable

### Consent for publication

Not applicable

### Availability of data and material

Sequence raw reads were deposited on NCBI SRA under BioProject PRJNA884767 and scripts used to analyze the data presented in this study can be obtained from https://github.com/LauRolon/NESARE.

### Competing interests

The authors declare that they have no competing interests.

### Funding

This study was funded by the National Institute of Food and Agriculture, U.S. Department of Agriculture, through the Northeast Sustainable Agriculture Research and Education program under subaward number GNE19-215 and Hatch Appropriations under project number PEN04646 and accession number 1017568.

### Author’s contributions

**MLR:** conceptualization, experimental work, data analyses, visualization, writing-original draft, writing-review and editing, project administration, funding acquisition; **TCC:** experimental work, data analyses, writing-original draft, writing-review and editing; **KEK:** writing-review and editing; **RFR:** writing-review and editing; **JK**: conceptualization, writing-review and editing, supervision, project administration, funding acquisition.

## Acknowledgements

The authors would like to extend their appreciation to the three dairy processing facilities that participated in the study and to Dr. Martin Wiedmann at Cornell University for donating the *L. monocytogenes* strains used in the study.

